# Persistent effects of intramammary ceftiofur treatment on the gut microbiome and antibiotic resistance in dairy cattle

**DOI:** 10.1101/2023.07.17.549362

**Authors:** Karla A. Vasco, Samantha Carbonell, Rebekah E. Sloup, Bailey Bowcutt, Rita R. Colwell, Karlis Graubics, Ronald Erskine, Bo Norby, Pamela L. Ruegg, Lixin Zhang, Shannon D. Manning

## Abstract

Intramammary (IMM) ceftiofur treatment is commonly used in dairy farms to prevent mastitis, though its impact on the cattle gut microbiome and selection of antibiotic-resistant bacteria has not been elucidated. Herein, we enrolled 40 healthy dairy cows after lactation: 20 were treated with IMM ceftiofur (Spectramast®DC) and a non-antibiotic internal teat sealant (bismuth subnitrate) and 20 (controls) received only bismuth subnitrate. Fecal samples were collected before (day −1) and after treatment (weeks 1, 2, 3, 5, 7, and 9) for bacterial quantification and metagenomic next-generation sequencing. Overall, 90% and 24% of the 278 samples had Gram-negative bacteria with resistance to ampicillin and ceftiofur, respectively. Most of the cows treated with ceftiofur did not have an increase in the number of resistant bacteria; however, a subset (25%) shed higher levels of ceftiofur-resistant bacteria for up to 2 weeks post-treatment. At week 5, the antibiotic-treated cows had lower microbiome abundance and richness, whereas a greater abundance of genes encoding extended-spectrum β-lactamases (ESBLs), CfxA, ACI-1, and CMY, was observed at weeks 1, 5 and 9. Moreover, the contig and network analyses detected associations between β-lactam resistance genes and phages, mobile genetic elements, and specific genera. Commensal bacterial populations belonging to Bacteroidetes most often possessed ESBL genes followed by members of Enterobacteriaceae. This study highlights variable, persistent effects of IMM ceftiofur treatment on the gut microbiome and resistome in dairy cattle. Antibiotic-treated cattle had an increased abundance of specific taxa and genes encoding ESBL production that persisted for 9 weeks, while fecal shedding of ESBL-producing Enterobacteriaceae varied across animals. Together, these findings highlight the need for additional studies that identify factors linked to shedding levels and the dissemination and persistence of resistance determinants on dairy farms in different geographic locations.

## INTRODUCTION

Globally, multi-drug resistant (MDR) bacteria were estimated to cause 4.95 (3.62-6.57) million human deaths a year, with third-generation cephalosporin-resistant *Escherichia coli* and *Klebsiella pneumoniae* among the leading causes of MDR deaths worldwide [1]. These resistant bacterial populations are also considered to be the most concerning and economically impactful antimicrobial-resistant threats in the U.S. [2]. Enterobacteriaceae with resistance to third-generation cephalosporins carry genes encoding extended-spectrum β-lactamase (ESBL) production, which also confer resistance to penicillins and monobactams. Hence, use of third-generation cephalosporins to treat human and for livestock production may contribute to the emergence of ESBL-producing Enterobacteriaceae. In the U.S, less than 1% of all antibiotics used in livestock correspond to cephalosporins, with the majority of use (80%) occurring in cattle [3]. At present, two cephalosporins are approved for use in dairy cattle, namely cephapirin (a first generation cephalosporin) and ceftiofur (a third-generation cephalosporin) [3, 4]. Ceftiofur is approved for use only via the parenteral and intramammary route for therapeutic indications including mastitis, metritis, respiratory disease, and foot rot [4].

Mastitis, an infection of the mammary gland, is the disease with the highest incidence in dairy cattle [5]; hence, ∼90% of dairy farms use intramammary (IMM) β-lactam antibiotics during the dry-off period to treat and prevent mastitis [5–7]. More specifically, a study of 37 Wisconsin dairy farms reported ceftiofur to be the most common β-lactam antibiotic used intramammarily to treat clinical mastitis and for prophylactic dry-cow therapy [5]. Ceftiofur has bactericidal activity against both Gram-negative and Gram-positive bacterial populations, low toxicity potential, and efficient penetration of most body fluids. Consequently, β-lactams are also used to treat a variety of pathologies in humans such as septicemia, urinary tract infections, respiratory infections, meningitis, and peritonitis. Although cephalosporins like ceftiofur are mainly excreted in the urine (61-77%), they have been detected in the biliary system (∼30%) [8], ileum, and colon (20% of plasmatic concentration) [9]. However, the effects of IMM ceftiofur treatment on the fecal microbiome and resistome, or collection of antibiotic-resistance genes, have not yet been determined.

A prior study using mathematical modeling predicted that parenteral ceftiofur therapy would reduce the total concentration of *E. coli* in cattle, but would lead to an increase in the fraction of ESBL-resistant *E. coli* [10]. Despite this prediction, several prior studies have not observed a correlation between ceftiofur treatment and an increase in the emergence of ESBL-producing bacterial populations [9, 11, 12]. Although one study of cows receiving systemic ceftiofur treatment in early lactation observed an increase in the abundance of resistant Enterobacteriaceae for 7-8 days, the increase was temporary and was not observed at 29-35 days [13]. Similarly, in feedlot cattle, the combined treatment of chlortetracycline and ceftiofur was linked to an increase in the number of resistant *E. coli* and ESBL genes [14], suggesting co-selection of these antibiotic resistance genes (ARGs). Because of these prior associations, we conducted a longitudinal study of dairy cattle to determine how IMM ceftiofur treatment impacts the gut microbiome and abundance of antibiotic resistant bacterial populations through the dry period and early part of lactation.

## METHODS

### Study design and sampling scheme

The aim of this study was to assess the effects of IMM ceftiofur hydrochloride (CHCL) treatment on the gut microbiome of dairy cows at dry-off, the last milking before the dry period (**Figure 1**). The study was conducted in 2019 (June-November) at the Dairy Cattle Teaching and Research Center at Michigan State University, which contained ∼230 lactating dairy cows. Forty healthy Holstein cows were enrolled at dry-off if they met the following inclusion criteria: no antibiotic treatment during the last 90 days of lactation and a somatic cell count (SCC) of <150,000 cells/mL using the most recent Dairy Herd Improvement Association (DHIA) test. The cows were randomly assigned to one of two treatment groups. The antibiotic-treated group (*n* = 20) received 4 IMM infusions (1 per mammary gland) that each contained 500 mg ceftiofur (SpectramastDC®; Zoetis Animal Health) after the last milking and an internal IMM teat sealant containing bismuth subnitrate (Orbeseal®; Zoetis Animal Health). The control group received only the internal IMM teat sealant without the SpectramastDC®. Cows were randomly assigned to treatment group and were matched based on parity and monthly milk production.

**Figure 1.**
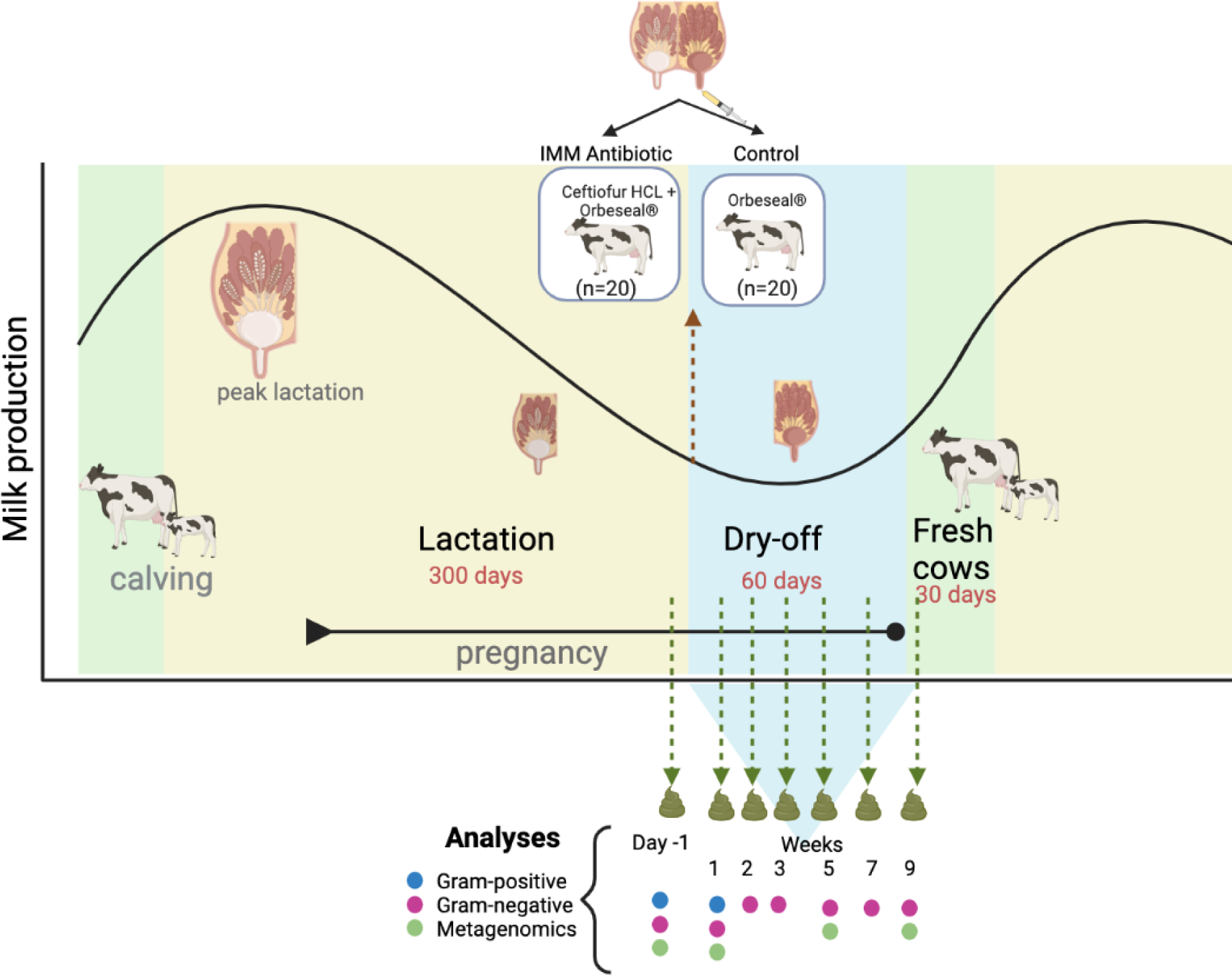
Study design showing the production stage and sampling time points for all 40 Holstein cows. Twenty dairy cattle received intramammary ceftiofur (IMM Antibiotic) and 20 matched dairy cattle receiving no IMM antibiotic treatment (Control). Cows were matched based on parity and monthly milk production at Day −1, which corresponds to the last day of lactation and the day prior to IMM treatment. Matched cows were sampled simultaneously following treatment at weeks 1, 2, 3, 5, and 7 during the dry-off period and again as fresh cows at week 9 (fresh). All cows remained healthy throughout the study. Figure created with BioRender.

Fecal grab samples were collected using clean obstetric sleeves on the last day of lactation, which corresponded to the day prior to IMM treatment (Day −1). The matched cows were re-sampled simultaneously at weeks 1, 2, 3, 5, and 7 during the dry-off period and again as fresh cows at week 9 (**Figure 1**). Each sample was homogenized by hand massage in a whirl-pak bag and immediately aliquoted for bacterial culture and DNA extraction for metagenomic next-generation sequencing (mNGS). For the latter, 0.25 g of feces per sample was preserved at −80°C in 750 ul of 190 Proof ethanol. Data about health status, ambient temperature, and diet were recorded at each time point for future analyses. Animals from both treatment groups were given the same diet formulation at each sampling, which corresponded to their physiological and productive stage at the time. All researchers were blinded to treatment status during sample collection and the subsequent laboratory analyses.

### Quantification of antibiotic-resistant bacteria

Total bacterial counts were quantified and presented as colony-forming units (CFUs) per gram (g) of feces. Moreover, the percentage of ceftiofur- and ampicillin-resistance was quantified for Gram-positive bacteria on day −1 and week 1 as well as Gram-negative bacteria on day-1 through week 9. Fecal samples were diluted at a concentration of 10^-1^ using 1 g of feces and 9 ml of 1X PBS and plated in duplicate on selective media using a spiral autoplater (Neutec Group Inc.). The media for Gram-negative bacteria was MacConkey lactose agar (MAC; Criterion®), whereas Columbia Nalidixic Acid agar (CNA; BD Difco ®) with 5% sheep blood was used for Gram-positive bacteria. Amphotericin B (4 μg/ml) was also added to inhibit fungal growth along with varying concentrations of antibiotics. For both Gram-negative and -positive bacteria, the ceftiofur (Cef) concentration was 8 μg/ml [15], while 32 μg/ml and 25 μg/ml of ampicillin (Amp) were used for Gram-negative and Gram-positive bacteria, respectively, per the Clinical and Laboratory Standards Institute (CLSI) guidelines [16]. The plates were incubated at 37°C for 24 hours under aerobic conditions (MAC) or in the presence of 5% carbon dioxide (CNA) (**Figure 2**). Media controls were plated to test each batch of MAC for the ability to inhibit Gram-positive bacteria with *Staphylococcus aureus* ATCC 29213 and *Enterococcus faecalis* ATCC 29212.

**Figure 2.**
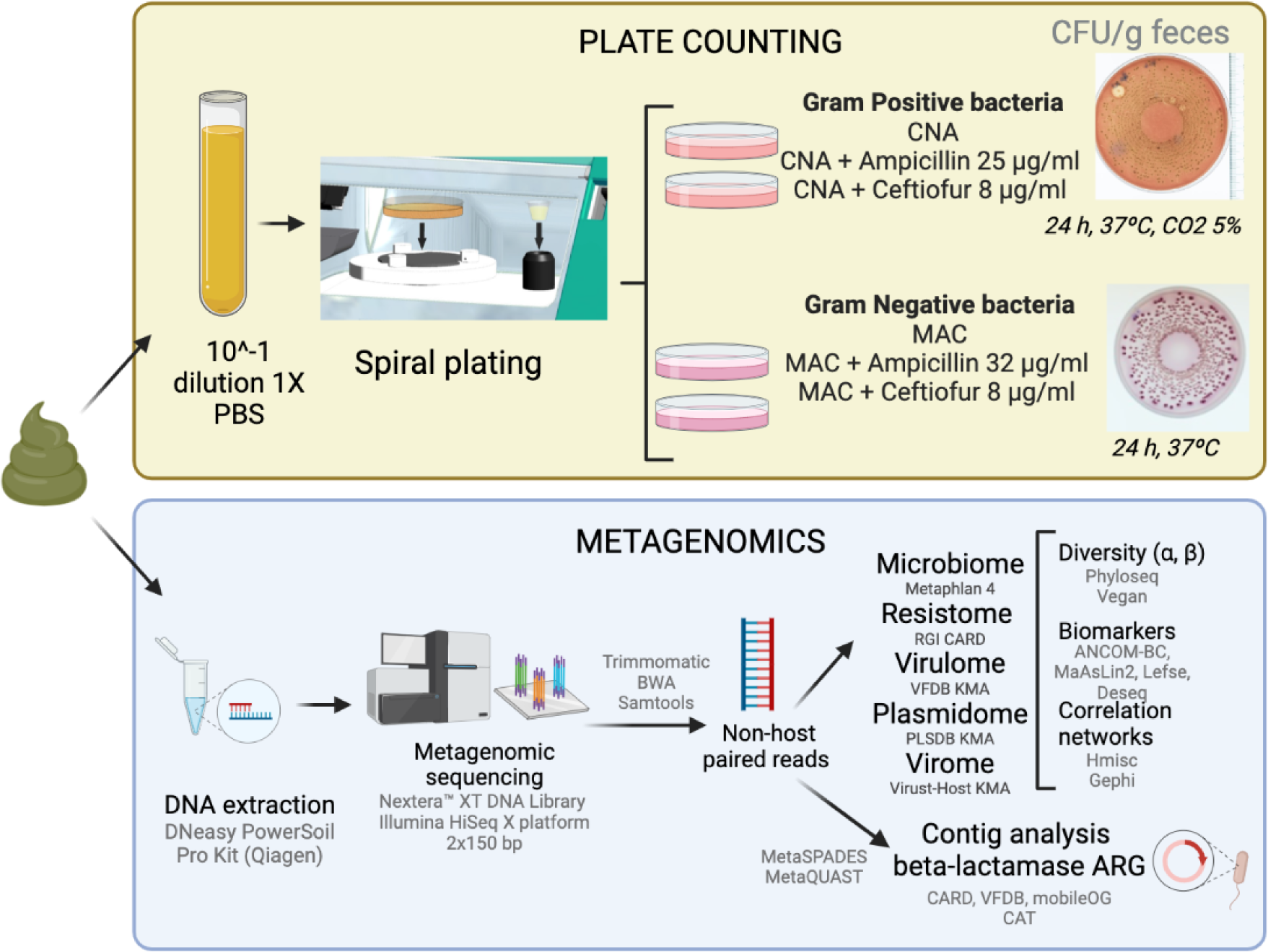
Summary of methods used for the quantification of Gram-positive and Gram-negative bacteria and metagenomics. The goal of these analyses was to identify the effects of intramammary (IMM) ceftiofur treatment on the cattle fecal microbiome using both culture-based methods and sequencing. Figure created with BioRender.com

The antibiotic concentration on MAC that inhibited susceptible (S) bacteria and enabled the growth of resistant (R) strains was tested with the following control strains: *E. coli* ATCC 25922 (Amp^S^, Cef^S^), *E. coli* ATCC 35218 (Amp^R^, Cef^S^), and three ESBL-producing *E. coli* strains (Amp^R^, Cef^R^) obtained from clinical samples in a prior study [17]. CNA media controls included ATCC 29212 (Amp^S^, Cef^R^), ATCC 29213 (Amp^S^, Cef^S^), *Listeria monocytogenes* ATCC 3382 (Amp^S^, Cef^R^), *L. monocytogenes* ATCC 19115 (Amp^S^, Cef^R^), *Streptococcus pneumoniae* ATCC 49619 (Amp^S^, Cef^S^), *Streptococcus equi* subsp. *zooepidemicus* ATCC 700400 (Amp^S^, Cef^S^), and *Streptococcus agalactiae* strain COH1 (Amp^S^, Cef^S^). Inhibition of Gram-negative bacteria was tested with *E. coli* ATCC 25922 and the ESBL-producing *E. coli* strains. Finally, biochemical identification of Gram-negative ceftiofur resistant strains was done with oxidase tests (OxiStrips^TM^, Hardy Diagnostics) and Chromocult® Coliform agar (Merck KGaA, Darmstadt, Germany) to test β-glucuronidase and β-galactosidase activity.

Paired non-parametric tests, Wilcoxon and Friedman, were used to compare the number of CFU/g and proportion of resistant bacteria between treatment groups and time points. These paired tests were necessary to account for repeated measures within each animal. In addition, linear mixed-effects models and subsequent analysis of variance (ANOVA) were applied to determine the impact of diet, days after treatment and ambient temperature (fixed effects) by controlling by individuals (random effect) in the bacterial counts with the R package nlme v.3.1-161 [18].

### DNA isolation and metagenomic next generation sequencing (mNGS)

Fecal DNA from samples collected on day −1 and weeks 1, 5, and 9, were selected for DNA extraction and sequencing. The samples were centrifuged at 16,000 rpm for 5 minutes at 4°C to remove the supernatant and residual ethanol, which was followed by two washes with 1 ml of molecular grade 1X PBS that was removed as done in the prior step. The DNeasy PowerSoil Pro Kit (Qiagen, Germantown, MD, USA) was used to extract DNA from each sample according to the manufacturer’s instruction followed by a wash step using the C3 solution to improve the DNA quality ratio (260/230). Genomic DNA was measured using a Qubit (1277.3 ng ± 310.5 ng of dsDNA) and sent to CosmosID (Rockville, MD, USA) for mNGS. Libraries were prepared with the Nextera™ XT DNA Library Preparation Kit (Illumina, San Diego, CA, USA) and sequenced on the Illumina HiSeq X platform 2×150 bp (**Figure 2**).

Paired-raw sequences were processed with Trimommatic v.0.39 [19] to remove low-quality reads and adapters used for Illumina sequencing. BWA v.0.7.15 [20] and Samtools v.1.4.1 [21] removed bovine DNA reads (*Bos taurus*, ARS-UCD1.2). Quality control check of the sequences was done with FastQC [22].

### Microbiome and resistome characterization

Non-host paired reads were analyzed using the Metaphlan 4 software [23] and the mpa_vJan21_CHOCOPhlAnSGB_202103 database to identify taxonomic features (**Figure 2**). The minimum read length and mapping quality value was set to 60 bp, and −1, respectively. The robust average quantile value was 0.1, while the Bowtie2 presets were “very-sensitive-local”. The normalized abundance score for each taxonomic feature was calculated by dividing the number of reads by the number of genome equivalents, which were determined by dividing the total number of base pairs by the estimated average genome size with MicrobeCensus [24].

The R package Phyloseq v.1.38 [25] was used to analyze alpha and beta diversity of the taxonomic profiling. The alpha diversity was calculated using the number of reads and measured with the Shannon index and Observed index, or richness. Paired Wilcoxon tests were used to compare alpha diversity estimates between treatment groups per time point, which accounts for repeated measures. Linear mixed-effects models were also used as described above for quantifying antibiotic-resistant bacteria. Comparisons were only made between treatment groups within each time point. Normalized abundances were used to calculate the beta diversity based on Bray-Curtis dissimilarities. Permutational multivariate analysis of variance (PERMANOVA) with 999 permutations and principal coordinate analyses (PcoA) were performed to compare the beta diversity between treatments and time points.

To characterize the resistome, the Resistance Gene Identifier (RGI) v.6.0.0 software [26] was used to analyze non-host paired metagenomic reads based on homology models. The Comprehensive Antibiotic Resistance Database (CARD) v.3.2.5 was aligned with RGI bwt using KMA with 20 bp k-mers as seeds. The settings specified the use of each query sequence to match only one template; results were reported at the drug class and allele levels. Resistance determinants based on SNP models, such as those identified with the rRNA, protein variant, and protein overexpression models were excluded. The depth of each ARG allele was normalized by dividing by the number of genome equivalents. Alpha and beta diversity were also measured.

Finally, to identify plasmids, virulence factors, and virus sequences, the PLSDB (updated on 06-23-2020) [27], VFDB setB (12-08-2022) [28], and Virus-Host (11-29-2022) [29] nucleotide databases were used. The k-mer aligner KMA v.1.4.3 [30] was employed with 20 bp k-mers, requiring each query sequence to match only one template. Normalization and diversity analyses were also performed.

### Biomarker identification

The analysis of differentially abundant features was carried out with three different approaches: 1) Linear Discriminant Analysis (LDA) Effect Size (LefSe), which identifies the effect relevance of a differential feature based on an algorithm that includes non-parametric tests and LDA [31]; 2) Analysis of compositions of microbiomes with bias correction (ANCOM-BC), which uses linear regression models and corrects for bias induced by sample differences [32]; and 3) Microbiome Multivariable Associations with Linear Models (MaAsLin2) [33] that uses generalized linear and mixed models. A consensus approach was used to ensure robust identification of differentially abundant features; only differentially abundant features (*P* < 0.05) identified with two or more of the three pipelines were reported.

### Characterization of β-lactamase carrying contigs

Metagenomic sequences were assembled using metaSPADES [34] and evaluated with MetaQuast [35]. The proportion of reads mapping a contig was identified with BWA v.0.7.15 [20] and Samtools v.1.4.1 [21]. Prodigal (PROkaryotic Dynamic programming Gene-finding Algorithm) [36] was used to translate contigs into amino acid sequences, which were then mapped to the protein databases CARD [26], VFDB [28], and mobileOG [37] using DIAMOND blastp [38] with a minimum sequence identity of 80%. Contigs carrying β-lactamases that confer resistance to cephalosporins were extracted with seqtk and taxonomically classified using the contig annotation tool (CAT) v.5.2.3 [39].

### Network analysis

Correlations between ARGs, plasmids, viruses, virulence factors, and bacterial genera were identified by calculating Spearman’s correlation coefficients; only coefficients (ρ) greater than 0.75 and p-values < 0.01 were included in the networks. Significant correlations were analyzed in R v.4.1.2. with the package Hmisc v.4.7-2 [40] and Gephi v.0.9.2 [41]. Network statistics including the degree and betweenness centrality were calculated in Gephi. The comparisons of centrality measures among β-lactam ARGs were analyzed between treatment groups and time points using non-parametric statistics.

## RESULTS

### Characteristics of the study population and sampling scheme

In this longitudinal study, 40 cows were enrolled at the end of lactation and had an average of 266.24 days in milk (DIM). Animals were matched based on parity and monthly milk production and pairs were randomly selected for the treatment or control group. No difference in the DIM was observed between the antibiotic-treated (*mean* = 262.69) and control (*mean* = 269.59) groups. Mastitis was ruled out in these cows as the somatic cell counts (SCC) in milk was an average of 34,8718 +/-23,602 cells/mL (antibiotic group *mean* = 35,300 cells/mL; control group *mean* = 34,4211 cells/mL). Cows received four diets that corresponded to their lactation phase including maintenance (day −1), dry (weeks 1-5), close-up (week 7), and fresh (week 9). All cows were pregnant and seemingly healthy during the study with most giving birth around the ninth week after dry-cow therapy. Fecal samples were collected from all animals through the 9-week period except for one cow in the antibiotic group. This cow had a C-section in week 9 and hence, a final sample was not obtained. Because sampling began in the summer and ended in the fall, the temperatures gradually decreased over the course of the study.

### Bacterial quantities and phenotypic resistance levels vary across samples and treatments

On average, the total number of Gram-negative bacterial colony forming units (CFUs) per gram of sample was 7.88 x10^5^ (±9.72 x10^5^), which was significantly lower in the ceftiofur-treated cows than the controls (*P* = 0.003). By contrast, the average total Gram-positive CFUs per gram of sample was 9.11×10^5^ (±1.16 x10^5^), which was not significantly different between treatment groups (*P* = 0.127).

Variation in total bacterial counts was also observed at different time points throughout the sampling period (**Figure 3A**). For instance, lower total Gram-negative bacterial counts were observed in the antibiotic-treated cows one week (*P* = 0.0148) and five weeks (*P* = 0.0487) following treatment, but not at weeks 2, 3, 7, or 9 (*P* > 0.05). For the Gram-positive bacteria, significantly higher counts were recovered in the control animals relative to the ceftiofur-treated animals one week after IMM treatment (*P* = 0.029; **Figure 3B**). Although a lower abundance of Gram-negative CFUs was observed one day before treatment (Day −1) in the antibiotic-treated (*mean* = 4.41×10^5^ CFUs) versus control cows (*mean* = 8.25×10^5^ CFUs), this difference was not significant (*P* = 0.057). No difference was observed in the Gram-positive bacterial counts between Day −1 and week 1 within treatment groups (*P* > 0.05) as well. Importantly, linear mixed-effects models revealed that the number of Gram-negative bacteria was not influenced by treatment (*P* = 0.32) but by changes in diet (*P* = 0.02), particularly in the amount of metabolizable energy (*P* = 0.002). The interaction between days after treatment and ambient temperature was also significant (*P* = 0.002), which was higher and lower in week 9, respectively. Such findings indicate that factors other than IMM ceftiofur treatment (e.g., diet, lactation phase, and temperature) impact the abundance of Gram-negative bacteria in cattle feces.

**Figure 3.**
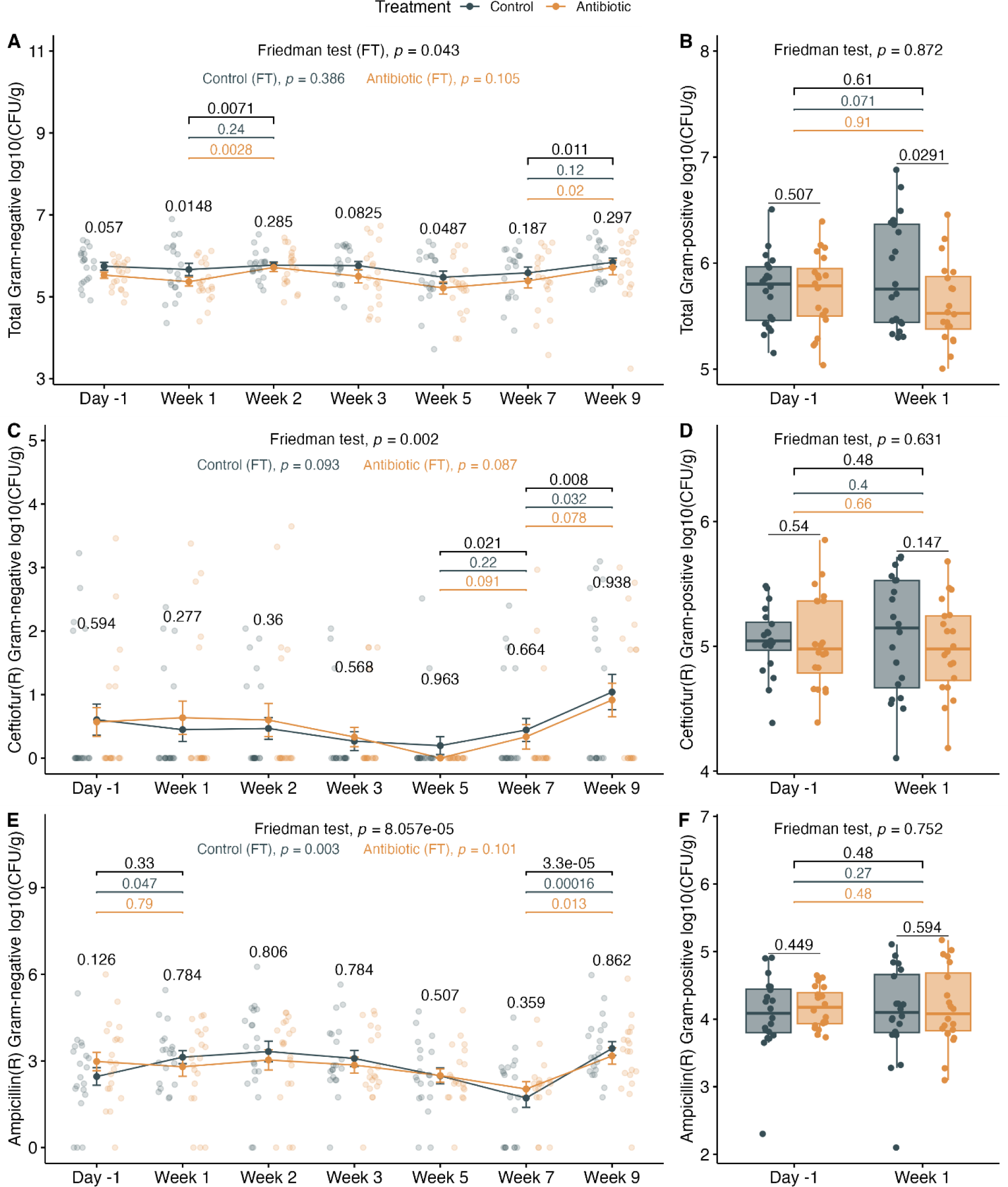
Number of bacterial colony-forming units (CFUs) per gram of feces. Total number (log_10_ CFUs/g) of **A**) Gram-negative bacteria; **B**) Gram-positive bacteria; ceftiofur resistant **C**) Gram-negative, and **D**) Gram-positive bacteria; and ampicillin-resistant **E**) Gram-negative and **F**) Gram-positive bacteria with (orange) and without (gray) intramammary ceftiofur treatment. Numbers are plotted before (Day −1) and after treatment for Gram-negative bacteria through 9 weeks and Gram-positive bacteria at 1 week. Line plots show means and standard error bars with dots showing sample counts. Boxplots indicate the median, lower, and upper quartiles, and the whiskers are extreme values in the distribution. P-values were calculated with a paired Wilcoxon test to compare treatment groups within a sampling point. The per animal variability over time was calculated with the Friedman test (FT), which is shown per treatment group for Gram-negative bacteria. Significant p-values between sampling points are shown for all animals (black), control (grey), and antibiotic-treated (orange) cows.

The number and percent of Gram-negative and Gram-positive bacterial populations with phenotypic resistance to ampicillin and ceftiofur were also determined. Notably, Gram-positive bacteria with resistance to both ampicillin and ceftiofur were present in 100% of the samples from both antibiotic-treated and control animals. In comparison, the percentage of samples with Gram-negative bacteria with resistance to ampicillin and ceftiofur was 90% and 24%, respectively. This difference, however, was not significant between the antibiotic-treated and control cows (*P* > 0.69).

Regardless of treatment status, significantly more Gram-negative bacteria were resistant to ampicillin (2.76%±10.60%) than ceftiofur (0.02%±0.09%) (*P* < 2.2e-16). Comparatively, a greater proportion of Gram-positive bacteria were resistant to ceftiofur (28.16% ±21.82%) compared to ampicillin (4.81% ± 6.06%) (*P* < 2.2e-16). Considerable differences were also observed in the percentage of Gram-negative bacteria with resistance to ampicillin (*P* = 0.001) and ceftiofur (*P* = 0.0015) across animals at the different time points (**Supplemental Figures S1A and S1B**). Among the samplings, the number of ampicillin resistant Gram-negative bacteria was significantly higher in the control group at weeks 1 (*P* = 0.041) and 2 (*P* = 0.021) compared to Day −1, but no differences were observed in the antibiotic-treated group (*P* > 0.3) (**Figure 3E**). Intriguingly, the total number of Gram-negative bacteria and the number of ampicillin resistant Gram-negative colonies increased at week 9 during pre-calving (*P* < 0.05) in both the treatment groups (**Figures 3C and 3E**). The number of Gram-negative bacteria with resistance to ceftiofur was also significantly higher in both groups at week 9 compared to weeks 5 and 7 (*P* <0.006), which was also true for Gram-negatives resistant to ampicillin (*P* <0.01). For the Gram-positive bacteria, the proportion with ampicillin resistance was also significantly higher in cows treated with ceftiofur (*P* = 0.0413) compared to controls (**Supplemental Figure S1D**). No difference, however, was observed in the quantity of Gram-positive CFUs with resistance to ceftiofur and ampicillin between treatment groups (**Figures 3D and 3F**).

### Metagenomic sequencing metrics

The metagenomic composition of cattle feces was analyzed for a total of 159 samples collected one day prior to treatment (day −1) and at weeks 1, 5, and 9 post-treatment. DNA extractions were performed in 13 batches by one individual using the same protocol. Fecal samples were randomly selected from both the ceftiofur-treated and control animals at the four time points for DNA extractions. The DNA library preparations and sequencing were performed in a single batch, except for one sample that was re-sequenced because of quality issues. The average number of reads (151 bp) per sample was 5.74 (±1.1) million, and no difference was observed in this number between treatment groups (*P* = 0.11) (**Table 1**). In week 5, however, samples from the antibiotic-treatment group had a lower number of reads (*P* = 0.035) (**Supplemental Figure S2**). The mean proportion of duplicate sequences was 11.36% (±2.25), while the GC content was 48.00% (±0.67) and 9.23% (±1.41) had failed sequences.

**Table 1.**
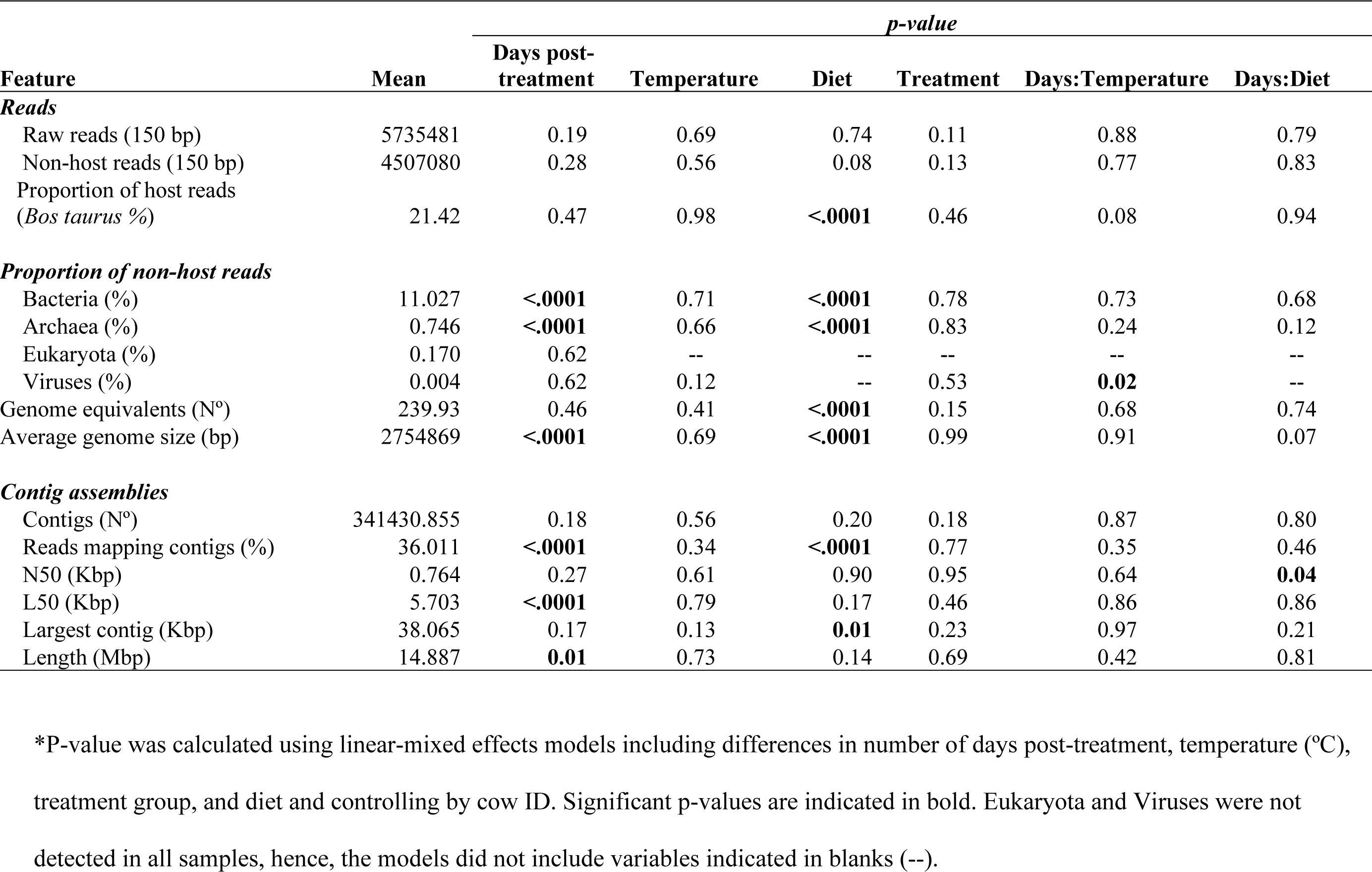
Metagenomic sequencing metrics from cattle fecal DNA.

After quality trimming, approximately 4,211.06 (±763.37) sequences were dropped per sample, corresponding to 0.07% (±0.01) of the raw reads. On average, 21.42% of the reads corresponded to bovine DNA. No differences were identified between treatments in the number of non-host paired reads (*P* = 0.129). However, cows treated with ceftiofur had a significantly lower number of non-host reads in week 5 (P = 0.041), but not in the number of genome equivalents (*P* = 0.062) (**Supplemental Figure S2**). The proportion of microbial taxa identified with Metaphlan 4 corresponded to 11.95% of the non-host reads, which varied significantly over time showing lower abundance in samples taken during the dry-off (weeks 1 – 5) and fresh (week 9) periods as compared to late lactation (day −1). The assembly’s length was also affected by the time (Table 1).

### Taxonomic profiling reveals differences across lactation phases

The microbiome was dominated by bacteria (92.29%), archaea (6.25%), eukaryotes (1.42%), and viruses (0.03%). The normalized abundance of microorganisms was significantly higher during the late lactation period (Day −1) compared to the dry-off and pre-calving periods (*P* < 0.001) (**Figure 4A).** Linear-mixed effects models were utilized to determine the contributing factors. In this analysis, lactation phase (*P* = 2.07e-07) and inclusion of a higher amount of grain in the diet prior to dry-off (*P* = 0.03) were associated with the observed differences in microbial abundance. Although environmental temperature did not significantly impact the taxonomic abundance (*P* = 0.87), differences in Shannon diversity were observed over the sampling period (*P* = 2.77e-10). The most diverse communities were detected before dry-cow therapy (Day −1) (**Figure 4B**). When comparing antibiotic-treated versus control cows, the abundance and alpha diversity of taxa were only significantly lower in week 5 (*P* = 0.01).

**Figure 4.**
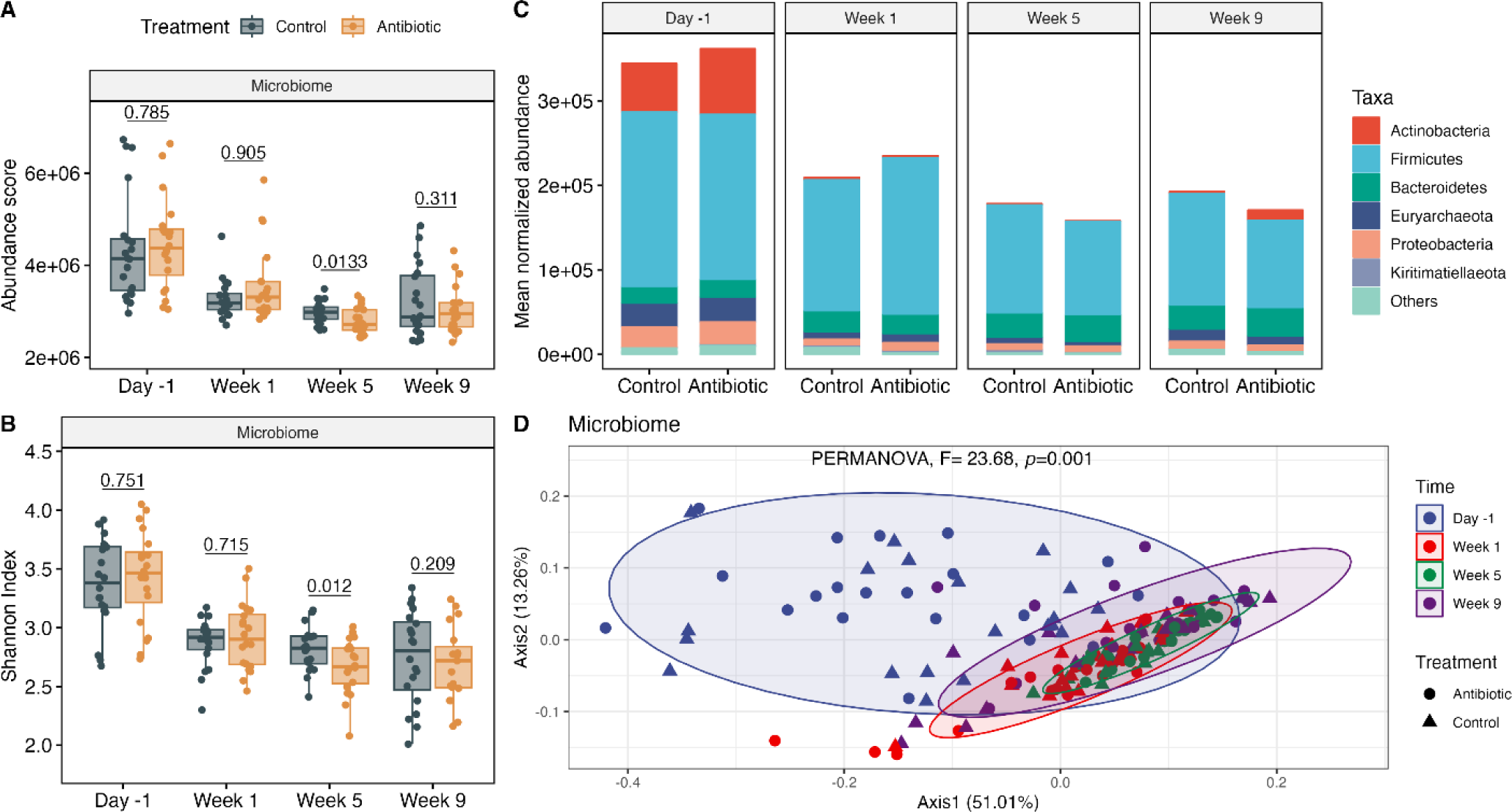
Microbiome diversity and composition before (Day −1) and 1, 5, and 9 weeks after dry-cow therapy. **A)** Normalized abundance and **B)** Shannon Index (alpha diversity) among ceftiofur-treated (orange) and control (grey) cows. Each boxplot shows the median, lower, and upper quartiles with the whiskers representing extreme values in the distribution. **C)** The mean normalized abundance of microbial taxa at the phylum level, and **D)** a PCoA of the Bray-Curtis dissimilarity. Ellipses in the PCoA are clustered by sampling point and contain at least 90% of the samples. P-values were calculated using a paired Wilcoxon test to compare treatment groups within a sampling point.

Significant changes in the bacterial composition were also detected over the sampling period as visualized in a relative abundance plot (**Figure 4C**) and a Bray-Curtis dissimilarity ordination (PERMANOVA, *F* = 23.68, *P* = 0.001) (**Figure 4D**). The day before dry-off (Day −1) was characterized by a higher abundance of Actinobacteria, Firmicutes, Euryarchaeota and Proteobacteria compared to weeks 1, 5, and 9. Stratifying by treatment status detected several differences in taxa abundance. Cows treated with ceftiofur, for instance, had a higher abundance of Ruminococcacea and a lower abundance of *Romboutsia* and Rickenellaceae one week after treatment compared to the control group (*P* < 0.05) (**Supplemental Figure S3**). At week 5, however, a lower abundance of several taxa including families Ruminococcacea, Lachnospiraceae, and Methanobacteriaceae, were detected in the antibiotic-treated cows.

Although a slight rebound in the Actinobacteria population was observed at week 9, this was only observed in the antibiotic-treated group (**Figure 4C**, **Supplemental Figure S3**). *Campylobacter* were also more abundant in the ceftiofur-treated cows compared to the controls. Overall, the differences in the taxonomic profiles between the treatment groups demonstrated a persistent effect of the antibiotic on certain bacterial groups, but not on the overall microbiome composition. No taxa were consistently affected over the 9-week period, though one taxon was persistently affected in weeks 1 and 9, three taxa in weeks 1 and 5, and five in weeks 5 and 9. Given that differences were observed due to time post-treatment, which included transitions in diet, temperature (high to low), and pregnancy stage, we only compared between treatment groups within each time point for microbiome abundance and diversity metrics.

### Resistome composition analyses identified persistent antibiotic resistance gene signatures

After treatment with ceftiofur, a significantly higher abundance of ARGs was observed in animals only in week 1 (*P* = 0.03) (**Figure 5A**), which was characterized by a lower Shannon index (*P* = 0.03) (**Figure 5B**). Similarly, the number of observed ARG alleles was lower in week 5 in cows treated with ceftiofur (*P* = 0.04). This finding indicates that IMM ceftiofur application resulted on a selective pressure for a group of antibiotic resistance genes in the intestinal environment in the short term. The main ARG drug classes identified were for resistance to tetracyclines (46.92%), followed by macrolides and streptogramins (19.04%), lincosamides (13.78%), and cephamycins (13.75%) (**Figure 5C**). The mean normalized allelic composition of ARGs varied significantly over time (PERMANOVA, *F* = 11.98, *P* = 0.001), though samples from different time points overlapped in the PCoA (**Figure 5D**). At the gene level, tetracycline resistance genes, *tet(W)*, *tet(Q)* and *tet(O)*, were the most abundant representing 28.49%, 10.28% and 6.12% of the ARGs detected, respectively. Other highly abundant genes were *mel* (9.09%), *cfxA2* (7.17%), *lnuC* (6.11%), and *bla*_OXA-608_ (5.04%) (**Supplemental Figure S4**).

**Figure 5.**
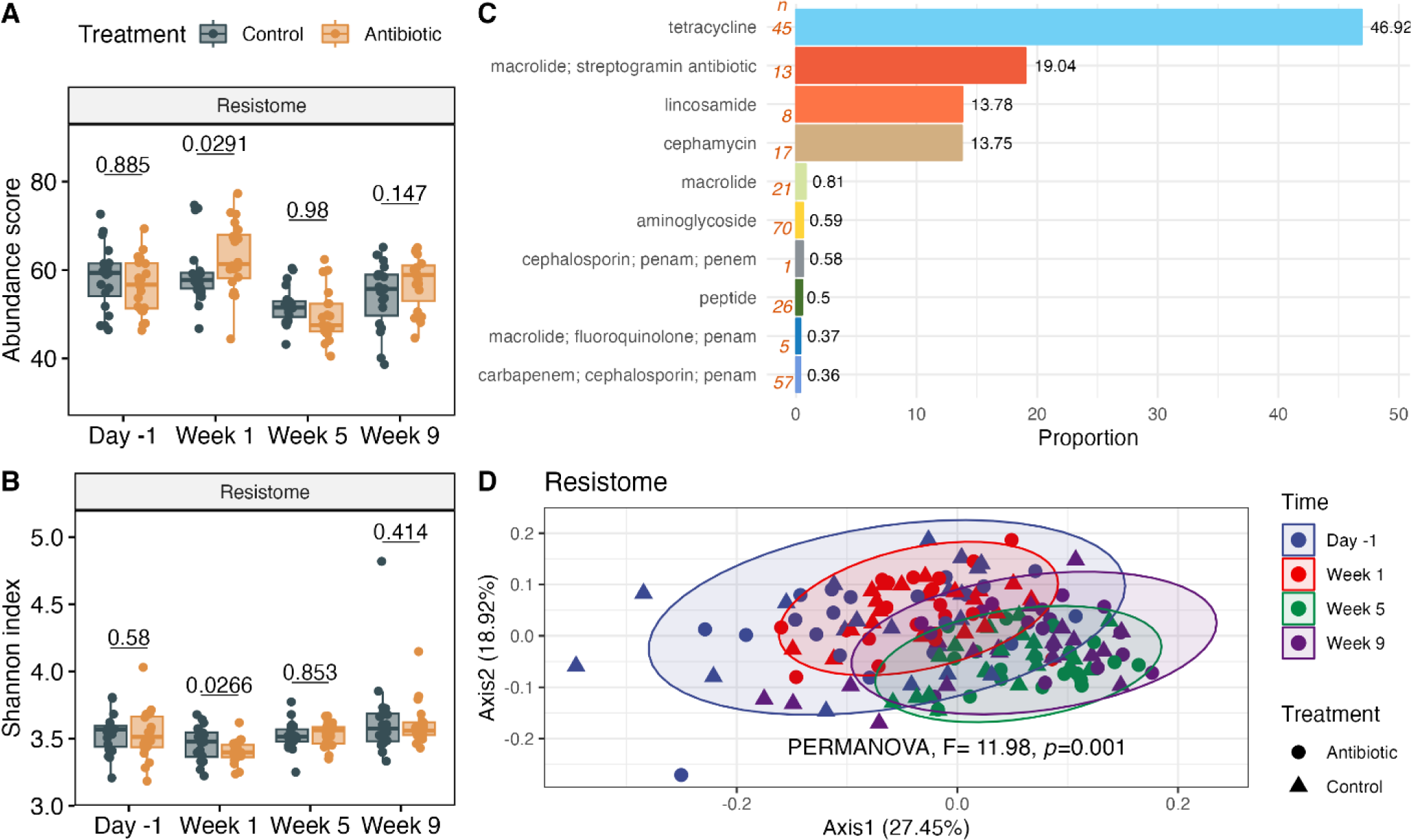
Fecal resistome composition of dairy cows during the 9-week study. **A**) The normalized abundance score and **B**) Shannon Index between antibiotic-treated (orange) and control (grey) animals. Each boxplot shows the median, lower, and upper quartiles with the whiskers representing extreme values in the distribution. **C**) The resistome composition at the drug class level showing the average proportion; “n” indicates the number of genes assigned to a given class. **D**) PCoA of the Bray-Curtis dissimilarity clustered by sampling point (ellipses contain at least 90% of the samples).

Importantly, a persistent increase in the abundance of genes encoding cephalosporin resistance was identified in weeks 1, 5 and 9 (*P* < 0.01) after IMM treatment with ceftiofur (**Figure 6A**). In comparison to the baseline measurement taken on Day −1, both treatment groups exhibited increased levels of cephalosporin resistance genes one week after IMM treatment (*P* = 7.7e-06). However, a significant difference was only observed in cows treated with ceftiofur (control, *P* = 0.062; antibiotic, *P* = 8.4e-05). Furthermore, the greatest abundance of cephalosporin resistance genes was found in week 9, which was significantly higher than in week 5 (control, *P* = 0.022; antibiotic, *P* = 0.0006). Genes important for cephalosporin resistance encoded antibiotic efflux pumps or were important for inactivation and reduced permeability to the drug. Antibiotic inactivation by β-lactamases was the main mechanism of resistance observed for the cephalosporins, showing a persistent increase in cows treated with IMM ceftiofur (**Figure 6B**). The β-lactamase (*bla*) genes encoding ESBL production, *aci1*, *cfxA2*, and *cfxA6*, were among those that increased over the sampling period (**Figure 6C**). Although controls also had these ESBL genes, more were identified only in the antibiotic-treated group at week 5, including *bla*_CMY-22_ and *bla*_CMY-59_. Additionally, co-selection of other ARGs including *qacG*, *tet(X4)*, and *arnA*, was observed in weeks 5 and 9 in the ceftiofur-treated cows (**Figure 6C**). These ARGs confer resistance to disinfectants, tetracycline, and peptides, respectively.

**Figure 6.**
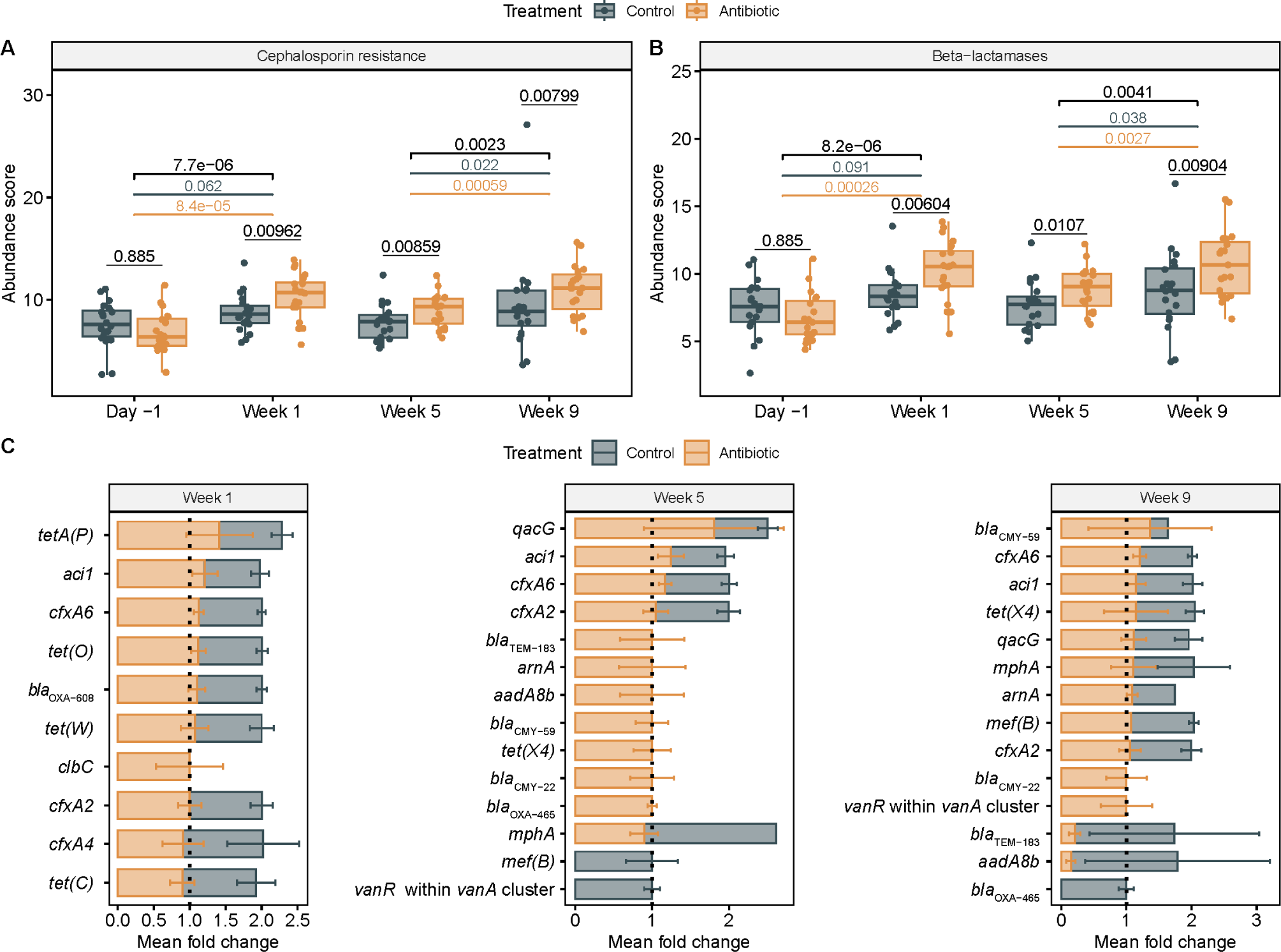
Effects of IMM ceftiofur treatment in the fecal resistome of cattle. Boxplots show the abundance score for genes encoding **A**) resistance to cephalosporins; and **B**) β-lactamases conferring resistance to cephalosporins. **C**) Differentially abundant ARGs identified after ceftiofur treatment; the mean fold change and standard error per treatment group is shown. The median, lower, and upper quartiles are shown in each boxplot with the whiskers representing extreme values in the distribution. P-values were calculated with paired Wilcoxon test to compare treatment groups within a sampling point. Significant p-values between sampling points are represented for all animals (black) as well as for control (grey) and antibiotic (orange) groups.

**Figure 7.**
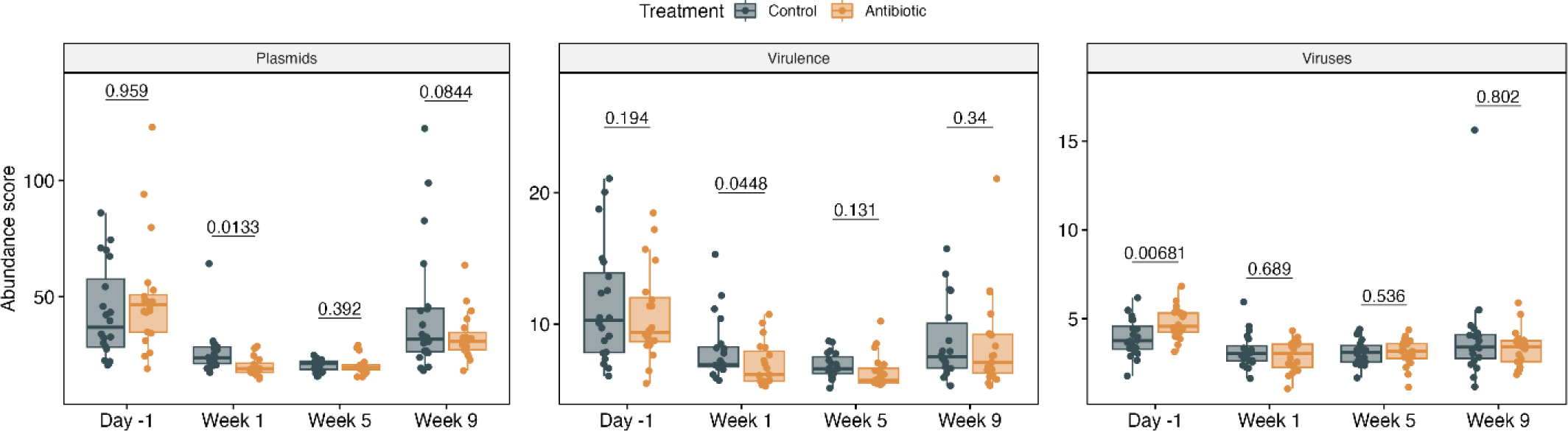
Effects of intramammary ceftiofur treatment on the abundance of plasmids, virulence factors and viruses. The median, lower, and upper quartiles are shown in each boxplot with the whiskers representing extreme values in the distribution. P-values were calculated with the Wilcoxon test to compare treatment groups within a sampling point.

### The plasmidome, virulome, and virome varied between treatment groups

The normalized abundance of plasmids and virulence genes was significantly lower in cows treated with IMM ceftiofur in the first week after treatment compared to the controls (*P* < 0.05), however, the number of observed features was similar between groups (*P* > 0.05). Despite the higher number of viruses identified prior to treatment in the antibiotic-treated group, no differences were observed at later time points. Nonetheless, there were significant differences observed in the mean plasmidome and virulome composition over time, with days −1 and week 9 as well as week 1 and week 5 forming two clusters in the PCoA (PERMANOVA, *P* = 0.001) (**Supplemental Figures S5A and S5B**). The virome composition also showed significant differences over time, but clear clusters were not observed in the PCoA (**Supplemental Figure S5C**). These analyses suggest that IMM ceftiofur lowered the abundance of plasmids and virulence factors in the short term.

### Multiple bacterial hosts had phenotypic or genotypic resistance to β-lactams

***Culture-based identification***. Among 882 Gram-negative bacterial isolates resistant to ceftiofur, 146 were preserved for further analyses; 72 were recovered from control cows and 74 from ceftiofur-treated cows. A maximum of 4 CFUs were selected per sample based on differences in morphology and lactose fermentation variation on MAC media. These colonies were recovered at day −1 (*n* = 26), week 1 (*n* = 25), week 2 (*n* = 17), week 3 (*n* = 17), week 5 (*n* = 5), week 7 (*n* = 10), week 9 (*n* = 44), and week 11 (*n* = 2). Biochemical assays classified 94 isolates as *E. coli*, 25 as other members of Enterobacteriaceae, and 27 as non-Enterobacteriaceae.

### β-lactamase-carrying contig (BCC) characterization

Among all 40 cows, 158 β-lactamase alleles (*bla*) conferring resistance to cephalosporins were identified in the fecal resistome. There were 792 contigs carrying these β-lactamase genes with an average size of 374.8 bp (±858.5 bp). The average coverage estimate was 2.3 (±8.9), while the number of 77 bp k-mers was 1269.54 (±8612.9). Co-localization of MGEs was identified in 318 (40.2%) of the contigs. β-lactamase genes *cfxA2*, *cfxA4*, and *cfxA6* were commonly found in those contigs containing MGEs for conjugation. These elements include *mobN*, *mobB*, *traC*, and *HMPREF1204_00020*, which encodes a DNA primase (EC2.7.7.-) that was linked to multidrug resistance in *Bacteroides* described in a prior study [42]. The gene encoding the SHV-160 β-lactamase was co-localized with the chaperonin gene, *groEL*, which is associated with plasmids and phages. Despite these findings, the taxonomic classification of the contigs with CAT was only possible for 20 contigs. These included genes encoding ACI-1 in Gammaproteobacteria, CfxA2, CfxA4, and CfxA6 in Bacteroidetes, EC-5 in *Treponema*, OXA-659 in Campylobacterales, SHV-160 in Proteobacteria and Bacteroidetes, and TEM-116 and TEM-183 in Enterobacteriaceae. For 18 of the 20 contigs, taxonomic assignments using CAT were based on a single ORF. Two contigs were exceptions: one classified as Aeromonadales was based on 2 ORFs, and another classified as *Bacteroides xylanisolvens* was based on 14 ORFs.

### RGI host assignations

The resistomes and variants database (CARD) also provided taxonomic assignations for various ARG alleles. Among the most abundant β-lactamases, the *cfxA2* sequences were assigned to *Phocaeicola* (45.94% of the allele reads), *Bacteroides* (38.8%), *Prevotella* (1.16%), *Parabacteroides* (9.65%), and *Butyricimonas* (4.44%). The *bla*_CFX-A6_ sequences were mostly assigned to uncultured organisms (97.55%) and *Bacteroides* (2.45%), whereas *aci1* was only assigned to *Acidaminococcus fermentans*. Genes encoding CMY-22 and CMY-59, which were detected only in the antibiotic-treated cows, were assigned to *E. coli* and *Klebsiella pneumoniae*, respectively. Other highly abundant β-lactamase genes were common in cows from both treatment groups including those encoding OXA-608, which was assigned to *Campylobacter jejuni*, and SHV-160 assigned to *Klebsiella pneumoniae*.

### Correlation networks

Correlations between β-lactamase genes and plasmids, phages, and virulence genes showed their potential ecological associations in the fecal microbiome. *E. coli* was the most common host of plasmids, phages and virulence factors correlated with β-lactamase genes, followed by *Klebsiella*, *Salmonella* and other Enterobacteriaceae (**Supplemental Figure S6**). However, the genera correlated with β-lactamase genes were primarily from phyla Firmicutes, Bacteroidetes, Actinobacteria, and Proteobacteria (**Supplemental Figure S7**). This discrepancy may be due to underrepresentation of non-Enterobacteriaceae sequences in the databases used to analyze accessory genes and associations with commensal bacteria. Moreover, no significant differences were observed in centrality measures between treatment groups at any time point, suggesting that the co-occurrence of β-lactamase genes with other genes and taxa was ecologically similar in both groups.

## DISCUSSION

It was estimated that ∼90% of dairy farms use IMM β-lactam antibiotics during the dry-off period to treat mastitis [5–7] despite the possibility of selecting for resistant bacterial populations. Of great concern is the emergence and selection of ESBL-producing Enterobacteriaceae, which are classified as a serious public health threat [1, 2]. Although the effect of IMM ceftiofur treatment has been studied in the milk microbiome, including five days with IMM 125 mg/day [43, 44] and a single application of 2 g of CHCL [45], the impact of this treatment on the gut microbiome had not been elucidated. Through this study, we have demonstrated persistent effects on the fecal microbiome due to a single 2 g dose of IMM ceftiofur via culture-based analyses and metagenomics. Relative to controls, antibiotic-treated cows had reduced microbial richness over time, differentially abundant taxa, and an increased abundance and persistence of β-lactam resistance genes that were associated with Enterobacteriaceae hosts and commensal bacteria. A subset of the ceftiofur-treated cows also had greater concentrations of ceftiofur resistant Gram-negative bacterial populations.

Following subcutaneous treatment, a prior study showed that Holstein steers had higher concentrations of CHCL in the gastrointestinal tract compared to ceftiofur crystalline-free acid (CCFA) [9], though only CCFA resulted in decreased fecal *E. coli* concentrations for up to two weeks. Similarly, parenteral ceftiofur treatment resulted in lower fecal *E. coli* concentrations for 3 days [12] and up to a month post-treatment [13] in two other studies. In the latter study of 96 dairy cows, systemic ceftiofur administration resulted in a significant increase in the level of ceftiofur-resistant Enterobacteriaceae, though the concentrations returned to baseline levels after one week [13]. Consistent with these findings, we observed a reduction in the total number of Gram-negative bacteria one week after IMM ceftiofur treatment. Enhanced recovery of Gram-negative bacteria with resistance to ceftiofur was observed for two weeks after the treatment. Re-emergence of ceftiofur resistance was also observed in the Gram-negative bacterial populations at 9 weeks (pre-calving) in both the antibiotic-treated and control animals, which is consistent with data generated in another study [13]. This increase was linked to sampling period and ambient temperature as well as diet, which increased the level of metabolizable energy given to fresh cows. Other factors that could contribute to the expansion of resistant Enterobacteriaceae populations include environmental acquisition of resistant strains, increased frequency of horizontal gene transfer, peri-parturient immune suppression, or increased contact with personnel. Regardless, it is important to note that *in vitro* bacterial quantifications do not distinguish between acquired and intrinsic antimicrobial resistance. Future studies should therefore focus on isolating the resistant strains for characterization using biochemical tests and whole-genome sequencing, which can define the genetic mechanisms of resistance as well.

Following IMM ceftiofur treatment, a lower abundance and diversity of taxa was detected in the fecal microbiome, which was also true for plasmids and virulence genes. Conversely, a higher abundance of ARGs was observed in the antibiotic-treated cows one week following IMM treatment. Because this difference was not observed in the subsequent time points, it suggests the temporary selection of resistant bacterial populations. Intriguingly, *Campylobacter* and *Bifidobacterium* were more abundant in the ceftiofur-treated cows as compared to controls, which is not surprising given that most *Campylobacter*, with the exception of *C. fetus*, have intrinsic resistance to third-generation cephalosporins [46]. In fact, nine β-lactamase genes were associated with *Campylobacter* including *bla_OXA-608_*, which was one of the most abundant ARGs detected.

Despite the temporary increase in ARG abundance and diversity observed one week after ceftiofur treatment, a subset of critically important genes persisted. Importantly, the antibiotic-treated cows had an exclusive and persistent increase in the abundance of ESBL genes (e.g., *aci1*, *cfxA,* and *bla*_CMY_) in the fecal resistome at each of the subsequent time points examined.

Although increases in the abundance of ESBL genes following parenteral ceftiofur treatment have been reported, no prior studies have examined the effect of IMM treatment. Steers receiving subcutaneous CCFA, for example, had a higher abundance of bacterial isolates harboring bla_CMY-2_ up to 4 days post-treatment, which resulted in co-selection of isolates containing *tet(A)* and *bla*_CMY-2_ after a subsequent chlortetracycline treatment for up to 26 days [14]. Similarly, Holstein cows treated with systemic CCFA had a higher abundance of genes encoding CfxA β-lactamases three days after treatment [47], while other studies reported an increase in *bla*_CMY-2_ in cattle feces for up to 10 days post-treatment when pure cultures were analyzed [12, 48].

Although the abundance of ESBL genes was higher in the ceftiofur-treated cows across the sampling period, an increase in cephalosporin-resistant bacterial populations (CFUs) was not observed. This discrepancy between the culture-based and sequencing methods could be attributed to the oxygenic environment and/or media used for cultivation. The hindgut microbiome is composed predominantly of anaerobic bacteria; thus, aerobic and microaerophilic conditions used for the quantification of Gram-negative and Gram-positive bacteria could only capture a fraction of the microbiota. Bacteroidetes members like *Prevotella* and *Bacteroides*, for example, are common Gram-negative anaerobes residing in the hindgut. Because these members were commonly found to carry genes encoding CfxA ESBLs [49], the resistant CFUs observed likely underestimate the actual levels of resistance, particularly given the high abundance of *cfxA* alleles detected. Likewise, *aci1* was the second most abundant ESBL gene and was previously reported in the Gram-negative Firmicutes *Acidaminococcus* [50] and Gram-positive genus *Bifidobacterium* [51]. These findings suggest that the increased abundance of ESBLs following IMM ceftiofur treatment were linked to changes in the abundance of anaerobic bacteria, which is consistent with our host analyses.

Indeed, identifying bacterial hosts and MGEs associated with β-lactam resistance genes in cattle feces is critical for developing new interventions, understanding the ecology of potential resistant threats that may emerge in farm environments, and defining risks associated with carriage of specific genes. As described herein, one approach to classify bacterial hosts is by identifying contigs or metagenome-assembled genomes containing genes encoding known β-lactamases. While culture identification of the resistant bacteria indicated 64% of the isolates were *E. coli*, metagenomic analyses showed that β-lactamase genes were mainly associated with commensal bacteria. A significant association was also identified between *bla*_CfxA_ and plasmid sequences, suggesting that horizontal gene transfer plays a key role in the acquisition of CfxA β-lactamase genes, particularly for members of phylum Bacteroidetes. Evidence of the relationship between Enterobacteriaceae and genes encoding the CMY, CTX, OXA, and TEM β-lactamase families was supported through the RGI analysis and co-occurrence networks showing correlations between these genes and plasmid sequences. Together, these results demonstrate the importance of horizontal gene transfer in the dissemination of antibiotic resistance within bacterial communities, particularly among members of the Bacteroidetes phylum and within the Enterobacteriaceae family.

Intriguingly, the abundance of Actinobacteria was significantly higher on day −1 compared to the subsequent time points. The most abundant family belonging to phylum Actinobacteria was Bifidobacteriaceae, which was represented mainly by the genus *Bifidobacterium*. Bifidobacteriaceae are implicated in the utilization of oligosaccharides in the colon resulting in the production of volatile fatty acids (VFAs) [52]. Differences in the composition of the fecal microbiome, primarily caused by the abundance of Actinobacteria observed on day −1 could be associated with differences in the diet. During late lactation, higher levels of dry matter intake and metabolizable energy as well as protein are consumed by cows compared to the dry off (weeks 1 – 7) and fresh (week 9) periods. However, further analyses of microbial metabolic pathways and metabolite composition are necessary to better explain how differentially abundant taxa may impact cow performance.

Although this study is the first to describe the impact of IMM ceftiofur treatment on the gut microbiome, it is important to highlight a few limitations. For instance, current resistome databases do not include all known ARGs from cattle samples and hence, novel resistance determinants may remain unclassified. Moreover, the identification of species and ARGs can be limited by a low number of metagenomic reads, as sequencing depth of ≥ 50 million reads is needed for complex microbial communities such as those residing in the bovine gut [53]. Since the proportion of microbial phyla and ARG classes was shown to be constant across various sequencing depths [53], we were able to detect the predominant and differential metagenomic features in this analysis. The shallow sequencing depth and short DNA segments (150 bp) examined, however, may have reduced our ability to accurately classify the bacterial hosts within each BCC since flanking regions are often not included. Such issues could have also contributed to the discordance observed between the sequence- and culture-based methodologies.

Consequently, future work involving use of third-generation sequencing platforms that sequence ultralong DNA segments such as the PacBio (40-70 kbp) or Oxford Nanopore Technologies (>100 kbp), is needed for confirmation and characterization of these regions [54]. Since the identification of differentially abundant features, including bacterial taxa and genes, tends to vary across bioinformatic pipelines, we applied three different approaches but only reported those features with significant p-values using at least two pipelines, as suggested previously [55]. Altogether, our analyses highlight those metagenomic features that are most impacted by IMM ceftiofur treatment.

## CONCLUSIONS

One application of IMM ceftiofur (2 g) at dry off contributed to an increase in the abundance of genes encoding resistance to cephalosporins and ESBLs in the fecal samples of antibiotic-treated cattle that persisted for nine weeks. Clinically important ESBL genes were mainly associated with Bacteroidetes and Enterobacteriaceae hosts as well as plasmid sequences, illustrating how ESBL-producing pathogens emerge and are selected for in this niche. While most of the cows given the prophylactic IMM ceftiofur treatment did not have altered microbiome compositions compared to the control cows, 25% had an increased level of ceftiofur-resistant Gram-negative bacteria for up to 2 weeks post-treatment. Indeed, the recovery of resistant CFUs was 14X greater in the antibiotic-treated versus control cows for up to two weeks after treatment. These findings demonstrate significant variation in the fecal shedding levels of cultivable bacterial populations across animals in this herd, which could be linked to selective factors such as diet, temperature, and lactation phase. Future studies should therefore focus on understanding the association between shedding and the dissemination and persistence of antibiotic resistance determinants in dairy farm environments across geographic locations.

## DECLARATIONS

### Ethics approval and consent to participate

The study protocol was approved by the Institutional Animal Care and Use Committee at Michigan State University (IACUC number ROTO201800166). The information and sample collections were approved by the Michigan State University (MSU) Dairy Cattle Teaching & Research Center in a written informed consent.

## Authors’ contributions

SM, PR, BN, LZ, & RE conceptualized the study and obtained funds for the project. PR organized and supervised the treatment assignments in the dairy farm and extracted epidemiological data. KV, SC, RS, & BB performed sample collection and bacterial culture experiments. KV performed DNA extractions, data analysis, interpretation, manuscript writing, and figure preparation. RC and KG carried out metagenomic sequencing via CosmosID. SM, PR, & LZ managed the project and supervised the study development. All authors read and approved the final manuscript.

## Acknowledgements

We present these findings in memory of Lorraine Sordillo-Gandy, MS, Ph.D. for her valuable contributions to the design and development of the project. We also thank the MSU Dairy Cattle Teaching & Research Center and Carmen Garcia, Zoe Hansen, Jose Rodrigues, Jaimie Strickland, Jennifer Brown, Jeffery Gandy, Robert West, and Aspen Robak, who participated in the field and/or laboratory work.

