## Supplemental figures for "Persistent effects of intramammary ceftiofur treatment on the gut microbiome and antibiotic resistance in dairy cattle"

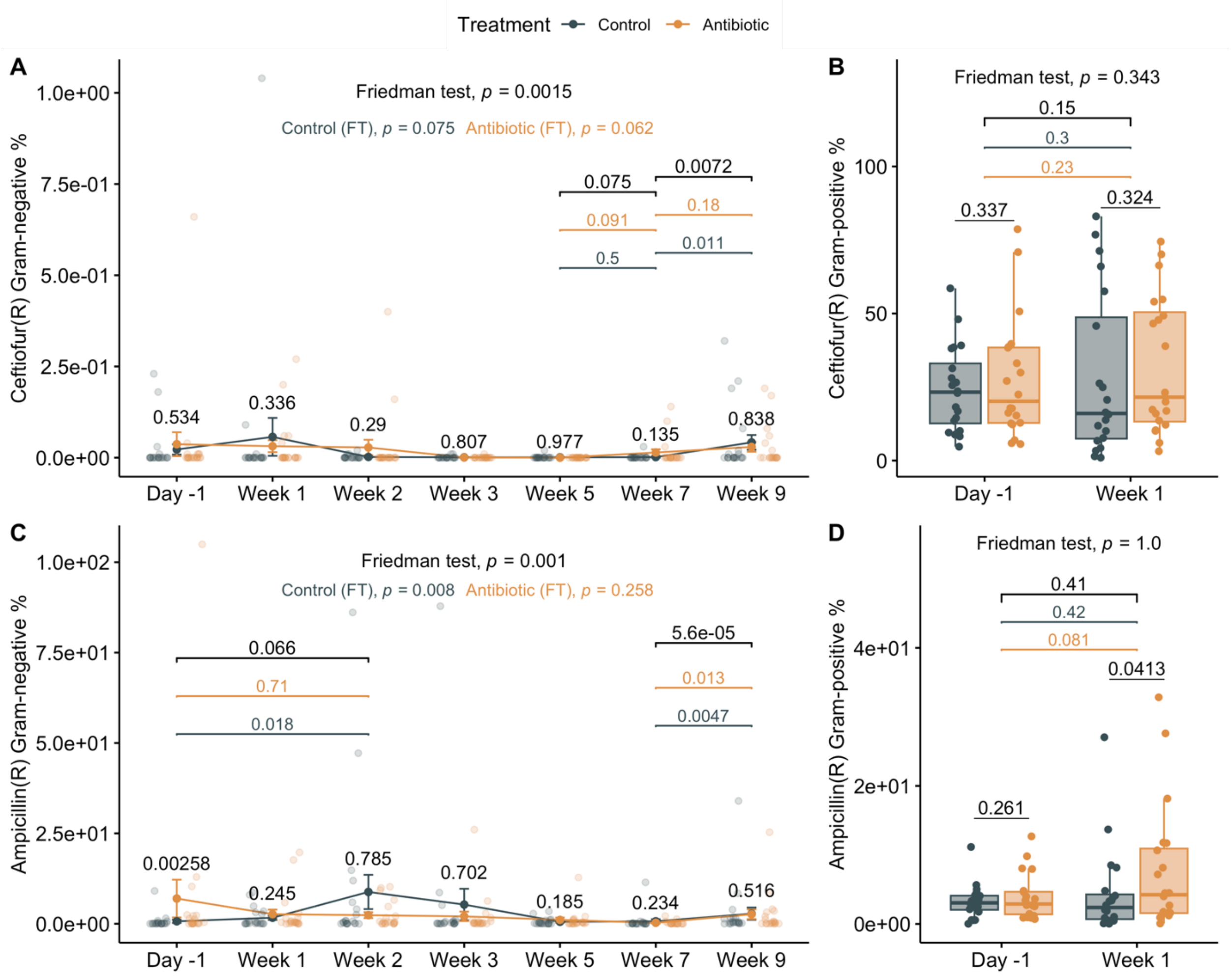


**Figure S1. Percentage of resistant bacteria to ceftiofur (A) Gram-negative and (B) Gram-positive; and resistant to ampicillin (C) Gram-negative and (D) Gram-positive.** The percentages were calculated based on the total number of CFU/g of feces. Line plots show means and standard error bars with sample counts represented as dots. P-values were calculated with paired Wilcoxon test to compare treatment groups within a sampling point. Boxplots indicate the median, lower and upper quartiles, and the whiskers represent extreme values in the distribution. The per animal variability over time was calculated with the Friedman test (FT), which is shown per treatment group for Gram-negatives. Significant p-values between sampling points are represented for all animals (black) as well as for control (grey) and antibiotic (orange) groups.

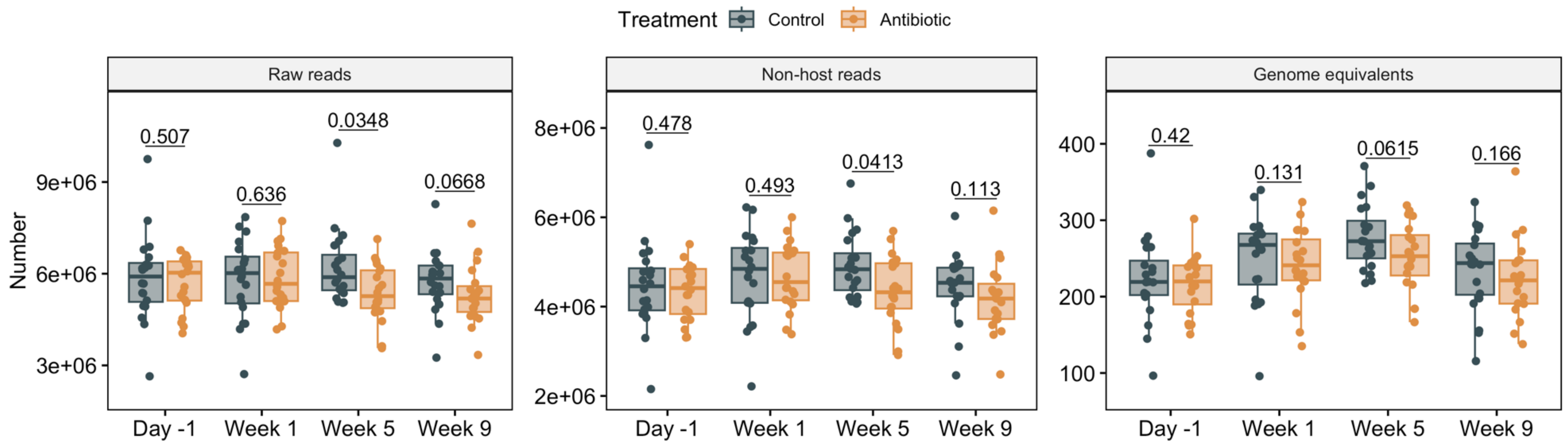


**Figure S2. Metagenomic sequencing metrics: raw reads, non-host reads and genome equivalents.** Boxplots indicate the median, lower and upper quartiles, and the whiskers represent extreme values in the distribution. P-values were calculated with paired Wilcoxon test to compare treatment groups within a sampling point. Genome equivalents were calculated with MicrobeCensus.

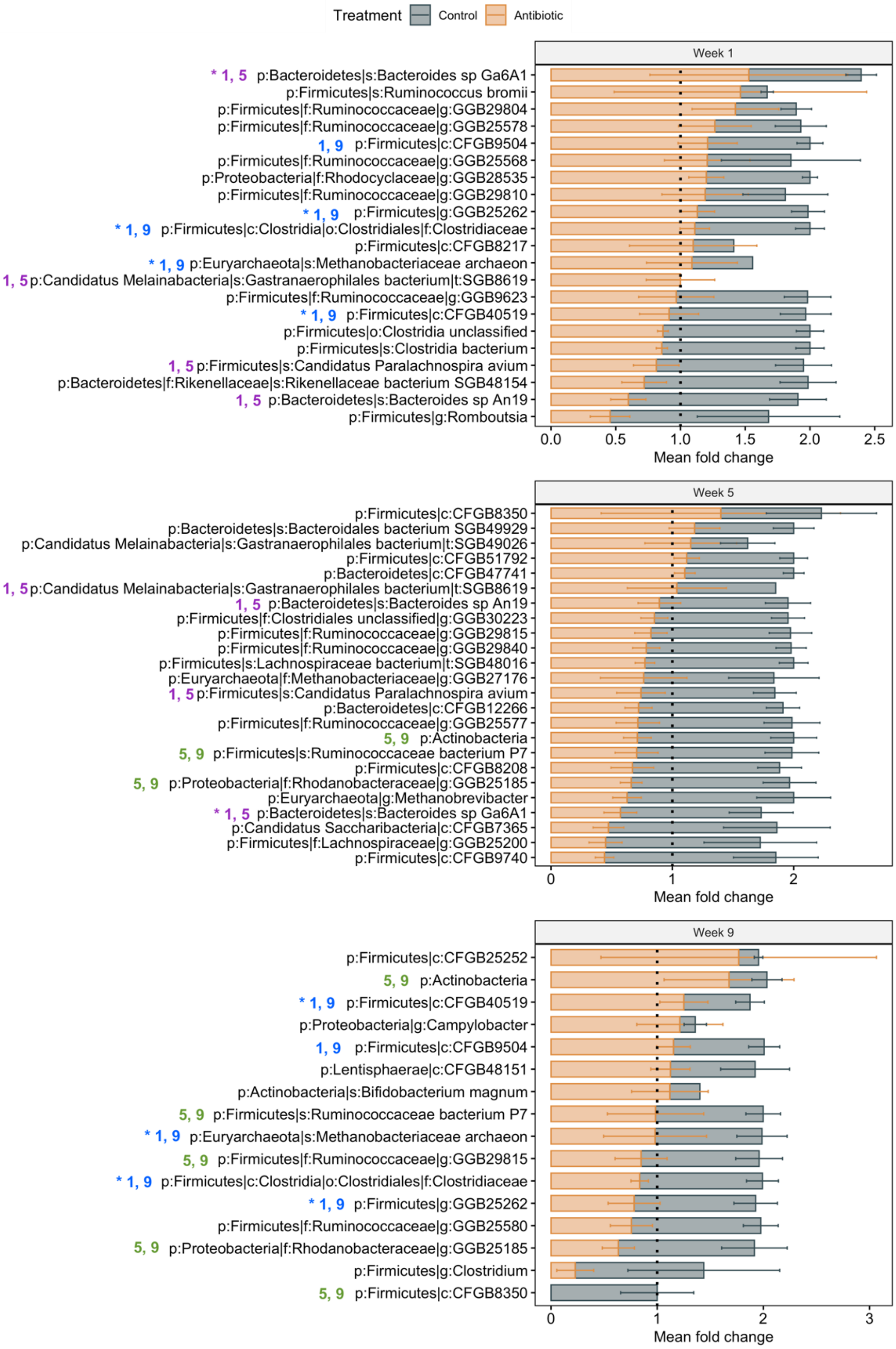


**Figure S3. Differentially abundant microbial taxa identified after IMM therapy indicating the mean fold change and standard error per treatment group.** Taxa that was significantly different in more than one point is indicated in blue for those observed in weeks 1 and 9, purple in weeks 1 and 6, and green in weeks 5 and 9. An asterisk indicate that a specific taxon was identified in two different time points but with opposite overrepresentation in a determined treatment group.

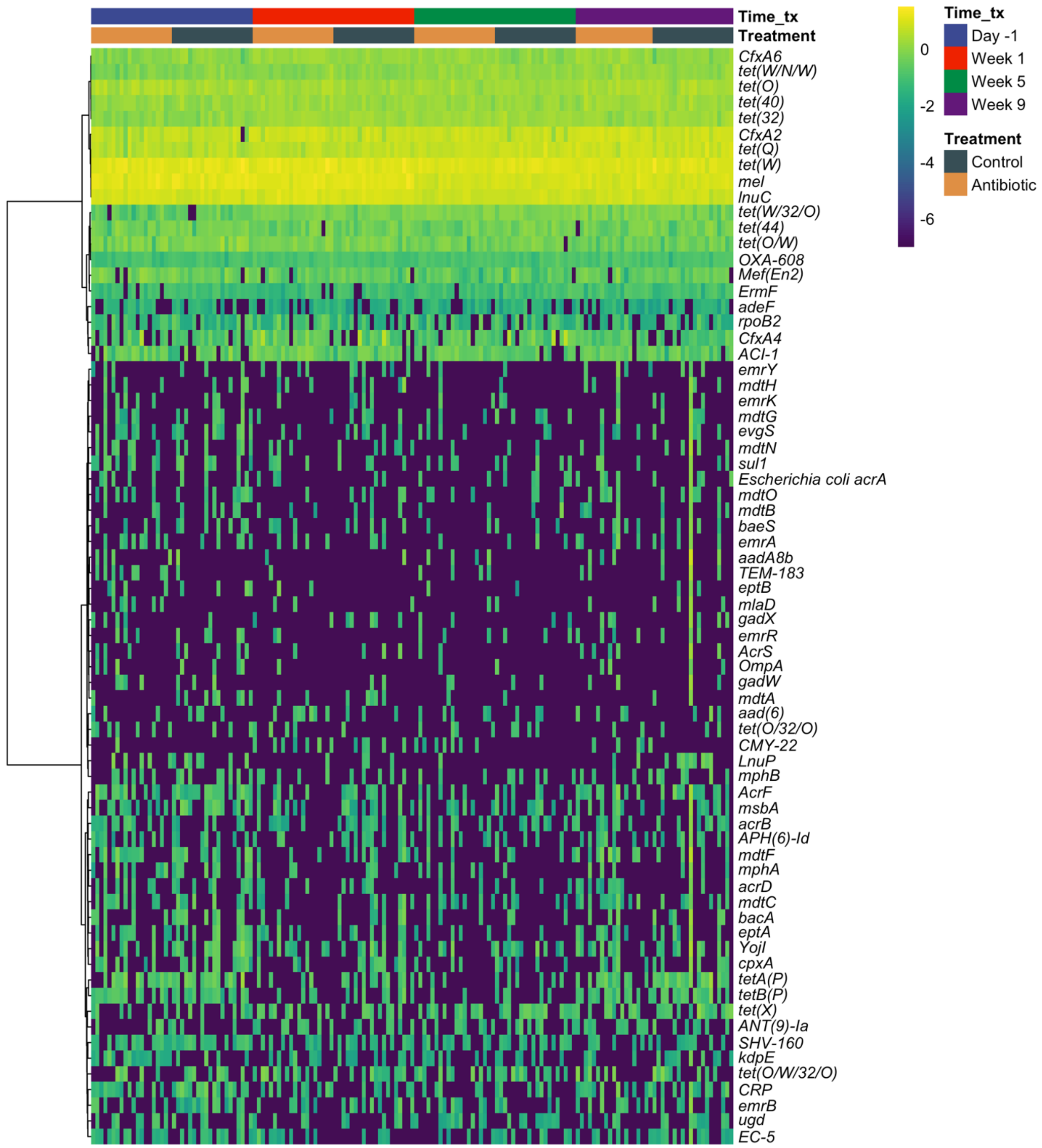


**Figure S4. Heat map of the 50 most abundant ARGs present in the fecal metagenome of cows.** Samples are organized by time point and treatment group. ARGs are clustered using the hierarchical clustering with the method Ward2. The values represent the logarithm 10 of the normalized abundance.

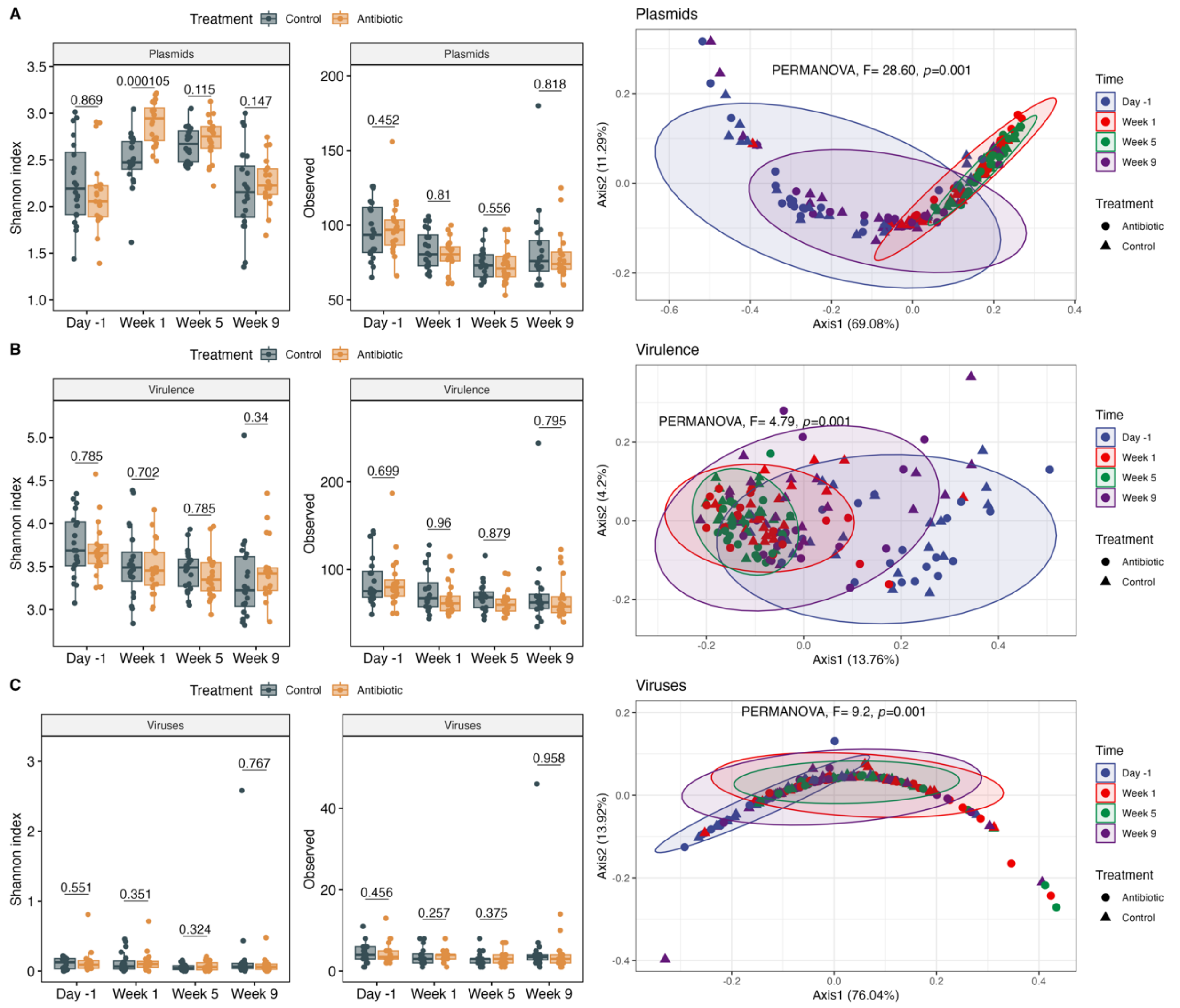


**Figure S5. Plasmid (A), virulence genes (B) and viruses (C) alpha and beta diversity in the fecal metagenome of dairy cows.** The Shannon index and observed features are indicated as boxplots that include the median, lower and upper quartiles, and the whiskers represent extreme values in the distribution. P-values were calculated with paired Wilcoxon test to compare treatment groups within a sampling point. PCoA of the Bray-Curtis dissimilarity is shown and clustered by time-point and treatment. Ellipses in the PCoA include 90% of the samples and PERMANOVA was calculated for differences between time points.

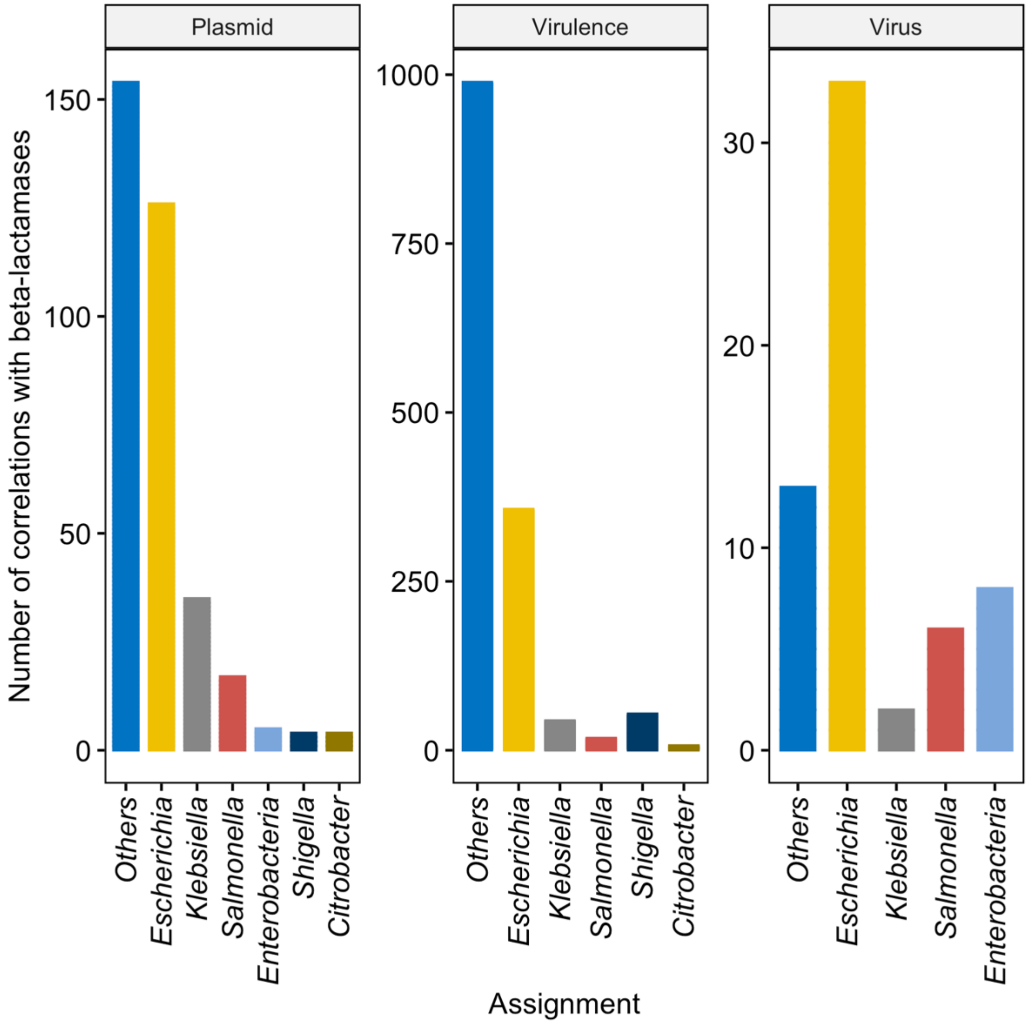


**Figure S6. Taxonomic assignment of plasmids, virulence factors and viruses correlated with β-lactamases.** The number of correlations identified between β-lactamases conferring resistance to cephalosporins are represented in the Y axis. Assignations were summarized at the genus level. The network included correlations ≥ 0.75 (*P* <0.01) calculated in a matrix with the normalized abundances of all samples (*n* = 159).

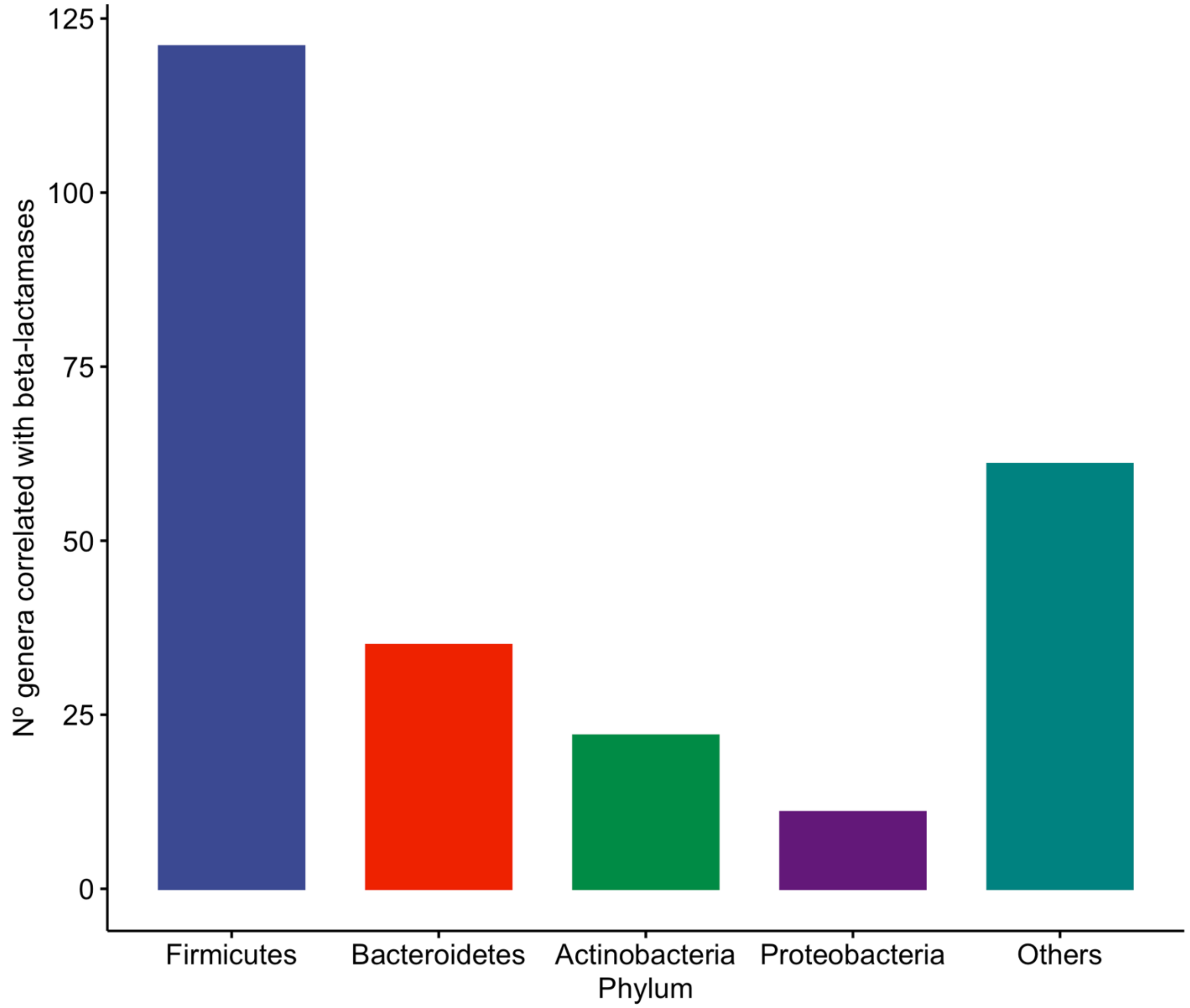


**Figure S7. Taxonomic assignment of genera clustered at the phylum level that was correlated with β-lactamases.** The number of correlations identified between β-lactamases conferring resistance to cephalosporins are represented in the Y axis. The network included correlations $\geq 0.75$ ($P < 0.01$) calculated in a matrix with the normalized abundances of all samples ($n = 159$).
